# A FRET Ligation Assay using Fluorescent Proteins for Bacterial Sortase Enzymes

**DOI:** 10.64898/2026.08.21.746329

**Authors:** Adam Wachsman, Erich G. Walkenhauer, Kaia Stover, Benjamin C. Richardson, Sophie N. Jackson, John M. Antos, Jeanine F. Amacher

## Abstract

Bacterial sortases are widely used in sortase-mediated ligation (SML) experiments for various protein engineering applications. The power of these enzymes to bind and cleave a specific recognition motif, followed by ligation to another substrate using a ping-pong reaction mechanism has numerous applications in vaccine and antibody/nanobody drug conjugate development, as a diagnostic and therapeutic tool, in creating novel insulin derivatives, etc. The most widely used sortase for SML is the class A sortase (SrtA) from *Staphylococcus aureus* (saSrtA), and its engineered derivatives. Despite its utility, saSrtA and other endogenous sortases are relatively inefficient enzymes and use can be limited by the need for specific recognition of the Cell Wall Sorting Signal (CWSS), sequence Leu-Pro-X-Thr-Gly, where X=any amino acid. Therefore, there is a need to continue to identify new tools for SML and to develop screening assays towards these endeavors. Here, we present optimization procedures for a FRET-based assay utilizing the GFP derivatives mTurquoise2 and SYFP2 to directly monitor formation of ligation products generated via SML. Similar to related assays, our recombinant substrates can be easily manipulated to screen either the substrate recognition motif, second substrate nucleophile, and/or sortase variants themselves. We believe continued optimization of this assay for a variety of high throughput uses in sortase screening strategies is possible, providing a proof-of-concept approach for continued SML reagent development.

## Introduction

Sortase-mediated ligation (SML) strategies are widely used in protein engineering for a variety of applications. Recent examples include creating multivalent SARS-CoV-2 vaccines and antibody/nanobody drug conjugates for cancer therapies, as a diagnostic and therapeutic tool for neurodegenerative disease, and in developing novel insulin derivatives, amongst many others.^1–13^ Sortase enzymes recognize a specific sequence, followed by targeted substrate cleavage, and ultimately ligation of two molecular building blocks to form the final product. This recognition involves only a small number of amino acid residues and the N- and C-terminal species outside of the sortase ligation site are not restricted to amino acids; therefore, the potential uses for SML are widespread.^11,14,15^ Although not exclusively, the vast majority of SML experiments utilize an engineered variant of the Class A sortase (SrtA) from *Staphylococcus aureus* (saSrtA), the first sortase enzyme identified over 25 years ago.^16,17^

SrtA enzymes are cysteine transpeptidases located on the surface of gram-positive bacteria, which are responsible for attaching substrates to the cell wall, e.g., proteins important for pathogenesis, environment sensing, etc.^18–20^ SaSrtA recognizes the Leu-Pro-X-Thr-Gly (where X=any amino acid) Cell Wall Sorting Signal (CWSS) sequence on substrates, for example, the virulence factor Staphylococcal Protein A (Spa), which interferes with the host immune response (CWSS = ^474^LPETG^478^).^21^ The catalytic efficiency of saSrtA is relatively low (*k*_cat_/*K*_m_ = 200 ± 30 M^-1^ s^-1^), based on previously reported values.^22^ Therefore, directed evolution was used to develop variants of saSrtA with increased catalytic efficiency (e.g., the pentamutant saSrtA5M with *k*_cat_/*K*_m_ = 23,000 ± 3,000 M^-1^ s^-1^) or to vary the recognition motif (e.g., SrtAβ which recognizes the LMVGG sequence, with *k*_cat_ = 0.018 s^-1^ and *K*_M_ = 0.128 mM, or *k*_cat_/*K*_M_ = 140.6 M^-1^ s^-1^).^13,22^ Additional techniques, including the creation of chimeric enzymes and consensus design to increase thermodynamic stability and activity, have also been used.^23,24^

Despite the aforementioned improvements in saSrtA function, there remains interest in generating additional SrtA variants that have desirable properties such as enhanced catalytic activity and altered substrate selectivities. To identify these variants, it is necessary to have robust assays for assessing relative activity. To date, a variety of methodologies have been described for this purpose. Examples include FRET-based assays consisting of peptide substrates labeled with an N-terminal 2-aminobenzoyl (Abz) fluorophore and C-terminal 2,4-dinitrophenyl (Dnp) quencher, analyses using gel electrophoresis (specifically, SDS-PAGE), HPLC and mass spectrometry-based assays, and others.^25^ Bioluminescence assays using split luciferase substrates and fluorescence assays using split-GFP or GFP derivatives (EGFP and cpVenus) have previously been developed as well.^26–29^ While useful, there are important considerations for weighing the relative advantages of each of these techniques. For example, fluorescence-based assays using peptides labeled with a fluorophore (e.g., Abz)/quencher (e.g., Dnp) pair typically measure the initial peptide cleavage step of the reaction, and do not assess formation of the ligation product. Alternate peptide substrates can be used to directly monitor formation of ligation products,^30–32^ but the overall workflow of solid-phase peptide synthesis and HPLC/mass spectrometry assays can be relatively slow and low throughput. A benefit of protein-based assays, e.g., using luciferase or GFP derivatives, is that protein substrates can be recombinantly expressed and purified and offer the option of genetic manipulation (e.g., sequence variation). Protein assays must be designed carefully, however, and in our own work attempting to reproduce a luciferase-based sortase activity assay we encountered solubility issues, which is a recognized problem with luciferase.^33^

Therefore, here, we sought to pursue two goals with respect to development of recombinant protein-based sortase assay systems. The first was to see if FRET-based GFP systems for monitoring sortase activity could be expanded to other donor/acceptor derivative pairs, and the second was to investigate optimization based on linker sequence. Towards the first question, we chose to use the well characterized fluorescent proteins mTurquoise2 and Super Yellow Fluorescent Protein 2 (SYFP2), which are known to be suitable for FRET assays, and could provide an alternative to EGFP and cpVenus previously used (**Figure 1**).^29,34,35^ We then increased our dynamic range by inserting a tryptophan zipper sequence into both substrates. We conclude that this is a usable platform for medium or high throughput screens investigating sortase substrates and/or SrtA variants themselves. Additional linker optimization could also be implemented. Taken together, there are growing opportunities to design and/or identify improved SrtA variants for SML, and we present a second protein-based FRET assay for future development.

**Figure 1.**
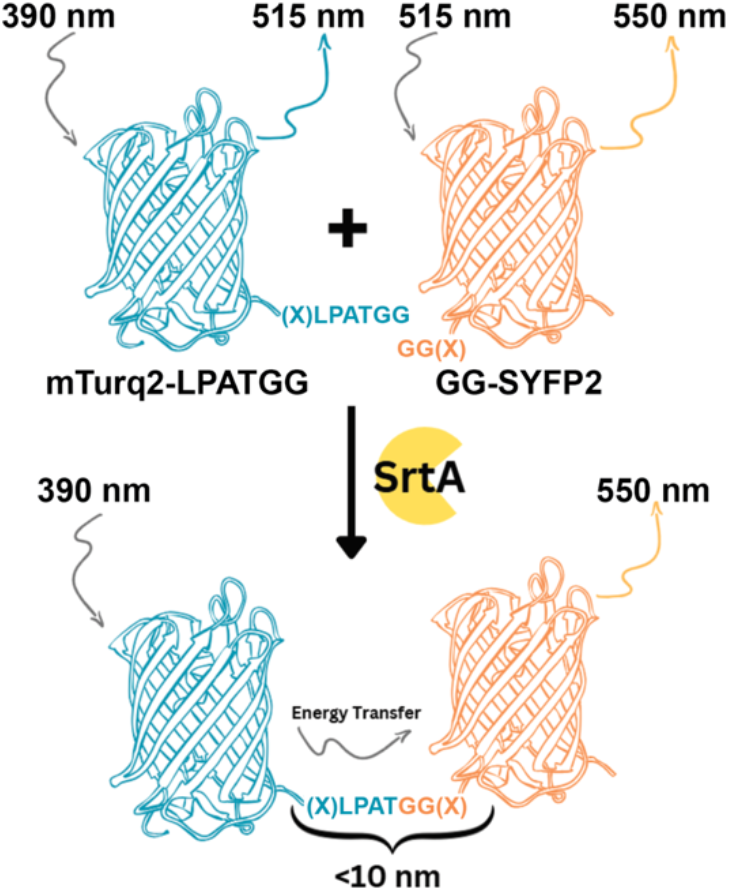
Structural schematic of the designed sortase-mediated ligation reaction between mTurq2 and SYFP2 fluorescent protein substrates. The mTurquoise2-LPATGG (mTurq2-LPATGG) and GG-SYFP2 fluorescent protein substrates are shown in cartoon representation. The wavelengths used for the ratio experiments are shown, with the excitation wavelengths indicated with gray arrows and emission wavelengths in blue (for mTurq2) and orange (for SYFP2).

## Materials and Methods

### Recombinant protein expression and purification

mTurquoise2 (mTurq2) and mTurq2_Trpzip), SYFP2 and SYFP2_Trpzip, EGFP-LPETGG, GG-cpVenus, and saSrtA5M were recombinantly expressed using *Escherichia coli* BL21 (DE3) cells in the pET28a(+) vector (Genscript).^36–38^ All sequences, with the exception of EGFP-LPETGG, contained an N-terminal 6x-histidine (6xHis) tag for purification, as well as a Tobacco Etch Virus (TEV) protease cleavage site (ENLYQS for mTurq2-LPATGG or ENLYQG for GG-SYFP2). EGFP-LPETGG contained a C-terminal 6xHis tag and no TEV protease cleavage site, based on the previously used sequence.^29^ All sequences are provided in the Supporting Information. Transformed cells were grown at 37 °C in LB media with 50 μg/mL kanamycin to an λ = 600 nm absorbance reading of 0.6-0.8, then induced using 0.15 mM IPTG for 18-20 hrs at 18 °C. Cells were harvested by centrifugation (6000 x *g* for 10 min at 4 °C), and resuspended in lysis buffer (0.05 M Tris pH 7.5, 0.15 M NaCl, and 0.5 mM EDTA) prior to sonication (Branson Ultrasonics Sonifier SFX250/SFX550 Cell Disruptor), and the whole-cell lysate was clarified by centrifugation (17,500 rpm for 30 min at 4 °C). The resulting supernatant was filtered through Miracloth (Sigma). Proteins were initially purified using a 5 mL HisTrap HP column (Cytiva) with wash (0.05 M Tris pH 7.5, 0.15 M NaCl, 0.02 M imidazole, and 0.001 M TCEP) and elution (wash buffer with 0.3 M imidazole) buffers.

After conducting immobilized metal affinity chromatography, TEV protease was added to the SYFP2 variants over 1.5 h at 34 °C at a ratio of ~1:100 (TEV:SYFP). A subtractive 5 mL HisTrap HP column was run using wash buffer, and the flow-through was collected. Following affinity chromatography, all proteins were further purified by size exclusion chromatography (SEC), using a HiLoad 16/600 Superdex 75 column (Cytiva) and SEC running buffer (0.05 M Tris pH 7.5, 0.15 M NaCl, and 0.001 M TCEP). The purity and monomeric state of purified proteins were assessed by SEC and SDS-PAGE. *S. aureus* sortase A pentamutnat (saSrtA5M) was expressed and purified as previously described, and following a similar protocol as described above.^36,37^ Proteins were then concentrated using an Amicon Ultra-15 Centrifugal Filter Unit (10,000 NWML), and aliquot concentrations were determined using theoretical extinction coefficients (280 nm) calculated using Benchling or ExPasy ProtParam: ε = 25900 M^-1^ cm^-1^ (mTurq2-LPATGG), 43890 M^-1^ cm^-1^ (mTurq2-LPATGG_Trpzip), 24870 M^-1^ cm^-1^ (GG-SYFP2), 41370 M^-1^ cm^-1^ (GG-SYFP2_Trpzip), and 15930 M^-1^ cm^-1^ (saSrtA5M).

### FRET ligation assays

FRET-based ligation assays were conducted in a black 96-well plate using a 100 uL reaction volume. Reactions contained 10% *(v/v)* 10x sortase reaction buffer (500 mM Tris pH 7.5, 1500 mM NaCl and 100 mM CaCl_2_). Final concentrations included: 10 μM EGFP-LPETGG or 50 μM mTurq2-LPATGG, 20 μM GG-cpVenus or 50 μM GG-SYFP2, and 5 μM saSrtA5M. Measurements were taken every 2 min for a duration of 2 h. For EGFP, cpVenus experiments, we used λ_excitation_ = 435 nm and λ_emission_ = 475 and 525 nm, as previously reported, although we ran our assays for t = 60 min (instead of 180 min, due to the observed activity) and at 25°C (not 37°C, as in the published validation experiment).^29^ For mTurq2, SYFP2 experiments, we used λ_excitation_ = 390 nm and λ_emission_ = 450 and 550 nm, and t = 120 min at 25°C. Each reaction was performed in triplicate. Data was analyzed and graphed using Excel and/or GraphPad Prism.

### SDS-PAGE analyses and quantification

A 100 μL SML reaction between mTurq2-LPATGG_Trpzip and GG-SYFP2_Trpzip proteins was performed *in vitro* using the conditions described above. Samples were taken after incubating for 0 and 2 h, then quenched using SDS sample buffer. Samples were diluted to 1:10 and 1:50, then analyzed using SDS-PAGE. An image of the resulting gel was uploaded to FIJI (Fiji Is Just ImageJ) for quantification. The 1:10 dilution produced the highest resolution bands and was therefore used for the quantification process. Individual reactant and product bands for t=0 and t=2 h timepoints were highlighted using the rectangle selection tool, and the gel analysis tool was used to represent the bands as curves. The area of each curve was then outlined and selected to produce a numerical value. Integrated values were compared to the total integrated density for each timepoint to determine relative protein quantities.

### AlphaFold3 modeling and structural analyses

AlphaFold3 was used to model substrate and/or products used.^39^ Structural analyses were performed using PyMOL.

## Results and Discussion

### Comparison to previous FRET fluorescent and peptide-based sortase cleavage assays

We first wanted to test if a previously established FRET protein assay could be used with additional fluorescent substrates. The previously developed FRET fluorescence assay used the GFP derivatives EGFP and cpVenus.^29,36,37,42^ We expressed and purified EGFP-LPETGG and GG-cpVenus substrates, based on the previously published sequence (EGFP-LPETGG) and a slightly modified GG-cpVenus substrate, where we incorporated an N-terminal 6x-His tag plus TEV protease cleavage site.^29^ Although we ran into fluorescence overload issues at the assay concentrations previously reported (100 μM EGFP-LPETGG and 200 μM GG-cpVenus), likely due to a higher reaction volume used, we successfully repeated the assay by reducing the relative substrate concentrations to 10 and 20 μM, respectively, with 1 μM enzyme (**Figure 2A**). As with the published data, we normalized our highest observed λ_em_525nm_/λ_em_475nm_ ratio to 100, such that all values were compared to the ratio of maximum intensity for the acceptor fluorophore divided by fluorescence at a wavelength where the donor fluorophore emission was minimized. Our results were consistent with previously reported data, wherein we observed a normalized dynamic range of 53-100 for saSrtA5M over the experiment time course, although we observed an earlier activity plateau than what was previously reported, likely due to the increased relative enzyme concentration (**Figure 2A**).^29^

**Figure 2.**
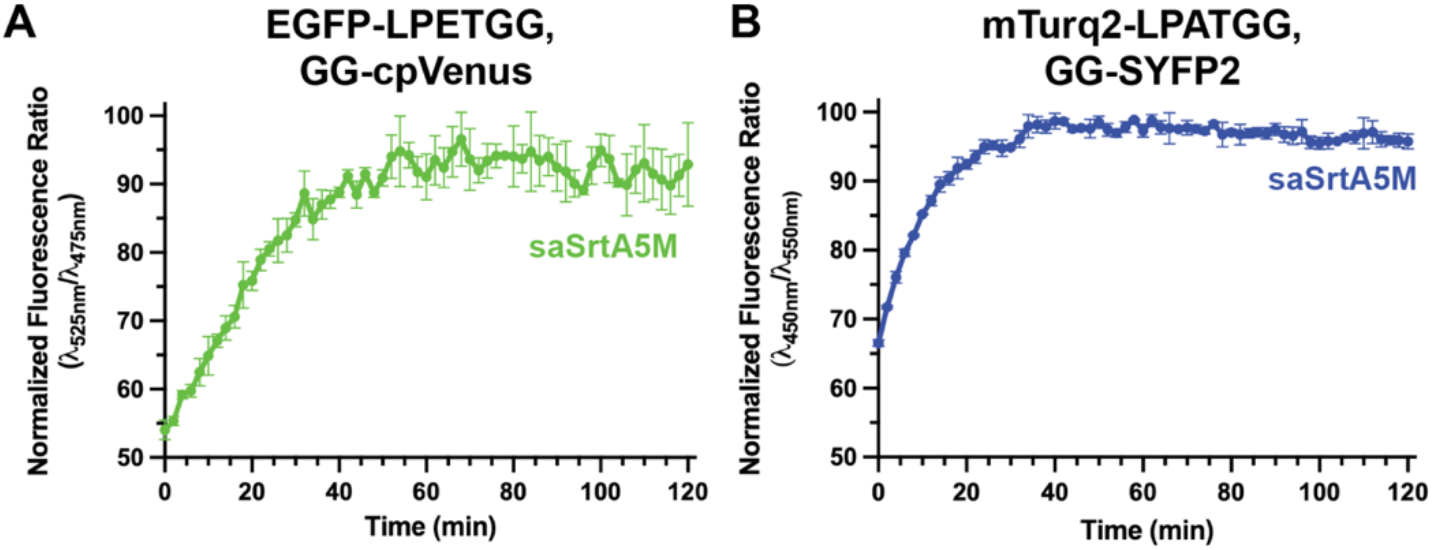
Comparison of different recombinant FRET protein pairs. (**A**) Validation of the previously published sortase-mediated ligation assay using the EGFP-LPETGG and GG-cpVenus substrate pair. Error bars are ± standard deviation from triplicate technical replicates. (**B**) Comparison assay for mTurq2-LPATGG and GG-SYFP2 substrate pair reported here. Error bars are ± standard deviation from triplicate technical replicates.

To test if we could use a different substrate pair in this assay, we downloaded the mTurquoise2 and SYFP2 fluorescent protein sequences, an established FRET pair, from FPbase.^40^ We appended a 6x His-tag for purification using immobilized metal affinity chromatography (IMAC) and a TEV protease cleavage site (ENLYFQ/S, cleavage indicated by the “/”) to the N-terminus of mTurquoise2. At its C-terminus, we included the sequence GGS followed by a sortase recognition motif (underlined): GGSLPATGGGGG (**Figure 1**). This substrate was termed mTurq2-LPATGG to reflect the addition of the sortase recognition motif. In our SYFP2 construct, we included an N-terminal 6x His-tag and TEV protease cleavage site (here, ENLYFQ/G) and termed this substrate GG-SYFP2. Following cleavage of GG-SYFP2 by TEV protease, this would generate an N-terminal sequence of G(GGGGS)_2_ followed by the native residues of SYFP2.^40^

We next wanted to test our mTurq2, SYFP2 substrates using a similar assay to the EGFP, cpVenus experiment. Here, our emission wavelengths were λ = 450 and 550 nm, based on fluorescence spectra of mTurq2 and SYFP2.^40^ We calculated λ_em_550nm_/λ_em_450nm_, and again normalized the highest ratio to 100 (**Figure 2B**). Although our normalized dynamic range (66-100) was not as high as for the EGFP-LPETGG, GG-cpVenus protein pair, we saw robust signal for an experiment with 50 μM mTurq2-LPATGG and 50 μM GG-SYFP2 using this approach (**Figure 2B**). Overall, this confirmed that our mTurq2, SYFP2 sortase-mediated FRET ligation assay was comparable to the previously developed EGFP, cpVenus version.

### An added “Trp zipper” increases FRET signal in sortase-mediated ligation assay

We next wanted to explore optimization of our relative fluorescence values. We reasoned that including a tryptophan zipper on both sides of the LPATG motif in the ligated product may orient the mTurq2 and SYFP2 chromophores for increased FRET efficiency. Tryptophan zippers are Trp-rich sequences typically engineered on either side of a β-hairpin, which can stabilize this conformation with large π-stacking energy.^50^ Although sequences can vary, many Trp zipper sequences are rich in alternating Trp-Thr (WTWT) residues.^50,51^ Notably, the use of Trp zipper sequences has been reported previously as a means for limiting the reversibility of SML reactions.^52^ Because we did not want to interfere with sortase recognition of our substrates, we engineered a WTWTW sequence N-terminal to the linker region in our mTurq2-LPATGG substrate (sequence: WTWTWGGSGGGGS). We also added the corresponding WTWTW sequence C-terminal to the repeating Gly residues in our GG-SYFP2 substrate (sequence: GGGGWTWTWGS). All full sequences are in the Supporting Information. The resulting substrates, termed mTurq2-LPATGG_Trpzip and GG-SYFP2_Trpzip, were expressed and purified like the other substrate variants.

Our Trpzip substrates showed increased signal in our FRET ligation assay; here, we observed an increased dynamic range (41-100), which was improved over the EGFP, cpVenus pair (**Figure 3A**). We also verified sortase-mediated ligation using SDS-PAGE and estimated ligation efficiency using the FIJI image analysis software (**Figures 3B, S1**). Based on this, our ligation efficiency was ~70% using saSrtA5M over a 2 h reaction period.

**Figure 3.**
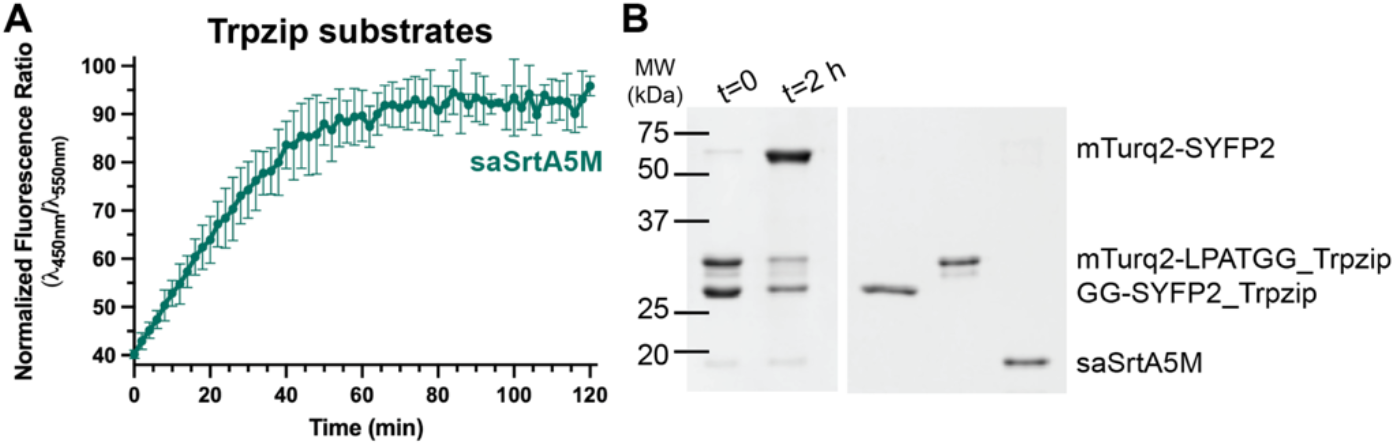
The dynamic range of the FRET ligation assay is improved by addition of a tryptophan zipper (Trpzip). (**A**) FRET ligation assay for mTurq2-LPATGG_Trpzip and GG-SYFP2_Trpzip substrate pair using normalized fluorescence ratio. Error bars are ± standard deviation from triplicate technical replicates. (**B**) SDS-PAGE analysis of the reaction at t=0 and t=2 h followed by quantification using FIJI software revealed an overall percent ligation for the Trpzip variants of ~70% using saSrtA5M. The FIJI integration data is in **Figure S1**.

## Supporting information

Supplemental Information for Wachsman et al.

## Concluding Remarks

As the options for sortase-mediated ligation experiments in protein engineering increase, the need for additional techniques to test relative activities also grows. While directed evolution has proven to be a powerful technique to identify saSrtA variants with greater catalytic efficiency,^22,24^ these approaches require a great deal of expertise and may not be accessible to all laboratories. Here, we present an additional assay using fluorescent proteins that can be utilized to probe SrtA and/or substrate variant sequences in a medium throughput manner. An advantage of this assay is that mTurquoise2 and SYFP2 can be recombinantly expressed and purified in relatively high yields and are stable proteins. The melting temperatures (T_m_) of mTurquoise2 (a derivative of cyan fluorescent protein) and SYFP2 (a derivative of yellow fluorescent protein) are likely 70-80°C; however, it should be straightforward to exchange these proteins with engineered thermostable versions, e.g., hyperfolder YFP (hfYFP, T_m_ = 94.2°C).^55^ These thermal stable substrates could then be used to identify thermal stable versions of SrtA enzymes, which could have increased uses for both *in vitro* and *in vivo* applications. While our assay is comparable to a similar assay using EGFP and cpVenus that was previously reported, we showed here that additional linker engineering (e.g., with the addition of tryptophan zipper sequences) can increase the overall dynamic range of the assay. Taken together, we describe an additional sortase-mediated ligation assay with potential for advancing this exciting field.

## Author Contributions

**Adam Wachsman**: Investigation (lead), Conceptualization (supporting), Writing – Review & Editing (supporting), **Erich G. Walkenhauer**: Investigation (supporting), Conceptualization (supporting), Formal analysis (supporting), Writing – Review & Editing (supporting), **Kaia Stover**: Investigation (supporting), **Benjamin C. Richardson**: Investigation (supporting), **Sophie N. Jackson**: Conceptualization (supporting), Writing – Review & Editing (supporting), **John M. Antos**: Formal Analysis (supporting), Supervision (supporting), Writing – Review & Editing (supporting), **Jeanine F. Amacher**: Conceptualization (lead), Visualization (supporting), Resources (lead), Funding Acquisition (lead), Supervision (lead), Writing – Original Draft Preparation (lead)

## Acknowledgements

The authors want to sincerely thank all members of the Amacher lab for research support and useful discussion. A special thank you as well to Dr. Peng Chen for helpful correspondence regarding their previously published assay. J.F. Amacher and J.M. Antos were funded by NIH 1R15GM154315-01. J.F. Amacher was additionally funded by a Cottrell Scholar Award from the Research Corporation for Science Advancement and a Henry Dreyfus Teacher-Scholar Award from the Camille and Henry Dreyfus Foundation. Additional funding was provided to E.G. Walkenhauer and A. Wachsman by Western Washington University Research and Sponsored Programs. A. Wachsman was also funded by NSF REU CHE-2243968. B. Richardson was also funded by a Jarvis Memorial Summer Research Award from the College of Science and Engineering at Western Washington University.

## References

1. MolinerMorro AJ, Sheward D, Karl V, Perez Vidakovics L, Murrell B, McInerney GM, Hanke L (2020) Picomolar SARS-CoV-2 Neutralization Using Multi-Arm PEG Nanobody Constructs. Biomolecules 10:1661.

2. Kwon S, Duarte JN, Li Z, Ling JJ, Cheneval O, Durek T, Schroeder CI, Craik DJ, Ploegh HL (2018) Targeted delivery of cyclotides via conjugation to a nanobody. ACS Chem. Biol. 13:2973–2980.

3. Beerli RR, Hell T, Merkel AS, Grawunder U (2015) Sortase EnzymeMediated Generation of SiteSpecifically Conjugated Antibody Drug Conjugates with High In Vitro and In Vivo Potency. PLoS One 10:e0131177.

4. Fu Z, Li S, Han S, Shi C, Zhang Y (2022) Antibody drug conjugate: the “biological missile” for targeted cancer therapy. Signal Transduct. Target. Ther. 7:93.

5. Woodham AW, Cheloha RW, Ling J, Rashidian M, Kolifrath SC, Mesyngier M, Duarte JN, Bader JM, Skeate JG, Da Silva DM, et al. (2018) Nanobody-Antigen Conjugates Elicit HPV-Specific Antitumor Immune Responses. Cancer Immunol Res 6:870–880.

6. Gébleux R, Briendl M, Grawunder U, Beerli RR (2019) Sortase A Enzyme-Mediated Generation of Site-Specifically Conjugated Antibody-Drug Conjugates. Methods Mol. Biol. 2012:1–13.

7. Li D, Martinez DR, Schäfer A, Chen H, Barr M, Sutherland LL, Lee E, Parks R, Mielke D, Edwards W, et al. (2022) Breadth of SARS-CoV-2 neutralization and protection induced by a nanoparticle vaccine. Nat. Commun. 13:6309.

8. Güttler T, Aksu M, Dickmanns A, Stegmann KM, Gregor K, Rees R, Taxer W, Rymarenko O, Schünemann J, Dienemann C, et al. (2021) Neutralization of SARS-CoV-2 by highly potent, hyperthermostable, and mutation-tolerant nanobodies. EMBO J. 40:e107985.

9. Hong H, Lin H, Li D, Gong L, Zhou K, Li Y, Yu H, Zhao K, Shi J, Zhou Z, et al. (2022) Chemoenzymatic Synthesis of a Rhamnose-Functionalized Bispecific Nanobody as a Bispecific Antibody Mimic for Cancer Immunotherapy. Angew. Chem. Int. Ed. 61:e202208773.

10. Obeng EM, Fulcher AJ, Wagstaff KM (2023) Harnessing sortase A transpeptidation for advanced targeted therapeutics and vaccine engineering. Biotechnol. Adv. 64:108108.

11. Amacher JF, Antos JM (2024) Sortases: structure, mechanism, and implications for protein engineering. Trends Biochem. Sci. 49:596–610.

12. Park C, Zhang Y, Jung JU, Buron LD, Lin N-P, Hoeg-Jensen T, Chou DH-C (2023) Antagonistic Insulin Derivative Suppresses Insulin-Induced Hypoglycemia. J. Med. Chem. 66:7516–7522.

13. Podracky CJ, An C, DeSousa A, Dorr BM, Walsh DM, Liu DR (2021) Laboratory evolution of a sortase enzyme that modifies amyloid-β protein. Nat. Chem. Biol. 17:317–325.

14. Antos JM, Truttmann MC, Ploegh HL (2016) Recent advances in sortase-catalyzed ligation methodology. Curr. Opin. Struct. Biol. 38:111–118.

15. Morgan HE, Turnbull WB, Webb ME (2022) Challenges in the use of sortase and other peptide ligases for site-specific protein modification. Chem. Soc. Rev. 51:4121–4145.

16. Ton-That H, Liu G, Mazmanian SK, Faull KF, Schneewind O (1999) Purification and characterization of sortase, the transpeptidase that cleaves surface proteins of Staphylococcus aureus at the LPXTG motif. Proc. Natl. Acad. Sci. USA 96:12424–12429.

17. Mazmanian SK, Liu G, Ton-That H, Schneewind O (1999) Staphylococcus aureus sortase, an enzyme that anchors surface proteins to the cell wall. Science 285:760–763.

18. Spirig T, Weiner EM, Clubb RT (2011) Sortase enzymes in Gram-positive bacteria. Mol. Microbiol. 82:1044–1059.

19. Mazmanian SK, Ton-That H, Su K, Schneewind O (2002) An iron-regulated sortase anchors a class of surface protein during Staphylococcus aureus pathogenesis. Proc. Natl. Acad. Sci. USA 99:2293–2298.

20. Jacobitz AW, Kattke MD, Wereszczynski J, Clubb RT (2017) Sortase transpeptidases: structural biology and catalytic mechanism. Adv. Protein Chem. Struct. Biol. 109:223–264.

21. Becker S, Frankel MB, Schneewind O, Missiakas D (2014) Release of protein A from the cell wall of Staphylococcus aureus. Proc. Natl. Acad. Sci. USA 111:1574–1579.

22. Chen I, Dorr BM, Liu DR (2011) A general strategy for the evolution of bond-forming enzymes using yeast display. Proc. Natl. Acad. Sci. USA 108:11399–11404.

23. Wójcik M, Vázquez Torres S, Quax WJ, Boersma YL (2019) Sortase mutants with improved protein thermostability and enzymatic activity obtained by consensus design. Protein Eng Des Sel 32:555–564.

24. Freund C, Schwarzer D (2021) Engineered sortases in peptide and protein chemistry. Chembiochem 22:1347–1356.

25. Popp MW-L, Antos JM, Ploegh HL (2009) Sitespecific protein labeling via sortase-mediated transpeptidation. Curr Protoc Protein Sci. 89:15.3.1-15.3.19.

26. Zhang C-H, Shao X-X, Wang X-B, Shou L-L, Liu Y-L, Xu Z-G, Guo Z-Y (2022) Development of a general bioluminescent activity assay for peptide ligases. FEBS J. 289:5241–5258.

27. Li N, Yu Z, Ji Q, Sun J, Liu X, Du M, Zhang W (2017) An enzyme-mediated protein-fragment complementation assay for substrate screening of sortase A. Biochem. Biophys. Res. Commun. 486:257–263.

28. Zhang J, Wang M, Tang R, Liu Y, Lei C, Huang Y, Nie Z, Yao S (2018) Transpeptidation-Mediated Assembly of Tripartite Split Green Fluorescent Protein for Label-Free Assay of Sortase Activity. Anal. Chem. 90:3245–3252.

29. Chen L, Cohen J, Song X, Zhao A, Ye Z, Feulner CJ, Doonan P, Somers W, Lin L, Chen PR (2016) Improved variants of SrtA for site-specific conjugation on antibodies and proteins with high efficiency. Sci. Rep. 6:31899.

30. Reed SA, Brzovic DA, Takasaki SS, Boyko KV, Antos JM (2020) Efficient Sortase-Mediated Ligation Using a Common C-Terminal Fusion Tag. Bioconjug. Chem. 31:1463–1473.

31. Li Y, Yang Y, Zhang C-Y (2018) Visualization and Quantification of Sortase Activity at the Single-Molecule Level via Transpeptidation-Directed Intramolecular Förster Resonance Energy Transfer. Anal. Chem. 90:13007–13012.

32. Kruger RG, Dostal P, McCafferty DG (2002) An economical and preparative orthogonal solid phase synthesis of fluorescein and rhodamine derivatized peptides: FRET substrates for the Staphylococcus aureus sortase SrtA transpeptidase reaction. Chem. Commun.:2092–2093.

33. Jones KA, Porterfield WB, Rathbun CM, McCutcheon DC, Paley MA, Prescher JA (2017) Orthogonal Luciferase-Luciferin Pairs for Bioluminescence Imaging. J. Am. Chem. Soc. 139:2351–2358.

34. Nolles A, Hooiveld E, Westphal AH, van Berkel WJH, Kleijn JM, Borst JW (2018) FRET Reveals the Formation and Exchange Dynamics of Protein-Containing Complex Coacervate Core Micelles. Langmuir 34:12083–12092.

35. Mastop M, Bindels DS, Shaner NC, Postma M, Gadella TWJ, Goedhart J (2017) Characterization of a spectrally diverse set of fluorescent proteins as FRET acceptors for mTurquoise2. Sci. Rep. 7:11999.

36. Piper IM, Struyvenberg SA, Valgardson JD, Johnson DA, Gao M, Johnston K, Svendsen JE, Kodama HM, Hvorecny KL, Antos JM, et al. (2021) Sequence variation in the β7-β8 loop of bacterial class A sortase enzymes alters substrate selectivity. J. Biol. Chem. 297:100981.

37. Kodama HM, Lindblom KM, Walkenhauer EG, Antos JM, Amacher JF (2024) Amino acid variability at W194 of Staphylococcus aureus sortase A alters nucleophile specificity. Protein Sci. 33:e5212.

38. Cox-Tigre N, Stewart ME, Tucker J, Walkenhauer EG, Wilce CS, Antos JM, Amacher JF (2026) Amino Acid Variants at the P94 Position in Staphylococcus aureus Class a Sortase Modulate Substrate Binding and Enzyme Activity. Biochemistry 65:1325–1339.

39. Abramson J, Adler J, Dunger J, Evans R, Green T, Pritzel A, Ronneberger O, Willmore L, Ballard AJ, Bambrick J, et al. (2024) Accurate structure prediction of biomolecular interactions with AlphaFold 3. Nature 630:493–500.

40. Lambert TJ (2019) FPbase: a community-editable fluorescent protein database. Nat. Methods 16:277–278.

41. Lau Y-TK, Baytshtok V, Howard TA, Fiala BM, Johnson JM, Carter LP, Baker D, Lima CD, Bahl CD (2018) Discovery and engineering of enhanced SUMO protease enzymes. J. Biol. Chem. 293:13224– 13233.

42. Gao M, Johnson DA, Piper IM, Kodama HM, Svendsen JE, Tahti E, Longshore-Neate F, Vogel B, Antos JM, Amacher JF (2022) Structural and biochemical analyses of selectivity determinants in chimeric Streptococcus Class A sortase enzymes. Protein Sci. 31:701–715.

43. Valgardson JD, Struyvenberg SA, Sailer ZR, Piper IM, Svendsen JE, Johnson DA, Vogel BA, Antos JM, Harms MJ, Amacher JF (2022) Comparative Analysis and Ancestral Sequence Reconstruction of Bacterial Sortase Family Proteins Generates Functional Ancestral Mutants with Different Sequence Specificities. Bacteria 1:121–135.

44. Johnson DA, Piper IM, Vogel BA, Jackson SN, Svendsen JE, Kodama HM, Lee DE, Lindblom KM, McCarty J, Antos JM, et al. (2022) Structures of Streptococcus pyogenes class A sortase in complex with substrate and product mimics provide key details of target recognition. J. Biol. Chem. 298:102446.

45. Vogel BA, Blount JM, Kodama HM, Goodwin-Rice NJ, Andaluz DJ, Jackson SN, Antos JM, Amacher JF (2024) A unique binding mode of P1’ Leu-containing target sequences for Streptococcus pyogenes sortase A results in alternative cleavage. RSC Chem. Biol. 5:30–40.

46. Schmohl L, Bierlmeier J, von Kügelgen N, Kurz L, Reis P, Barthels F, Mach P, Schutkowski M, Freund C, Schwarzer D (2017) Identification of sortase substrates by specificity profiling. Bioorg. Med. Chem. 25:5002–5007.

47. Hirakawa H, Ishikawa S, Nagamune T (2015) Ca2+ independent sortase-A exhibits high selective protein ligation activity in the cytoplasm of Escherichia coli. Biotechnol. J. 10:1487–1492.

48. Shrestha D, Jenei A, Nagy P, Vereb G, Szöllősi J (2015) Understanding FRET as a research tool for cellular studies. Int. J. Mol. Sci. 16:6718–6756.

49. Hirakawa H, Ishikawa S, Nagamune T (2012) Design of Ca2+-independent Staphylococcus aureus sortase A mutants. Biotechnol. Bioeng. 109:2955–2961.

50. Cochran AG, Skelton NJ, Starovasnik MA (2001) Tryptophan zippers: stable, monomeric beta-hairpins. Proc. Natl. Acad. Sci. USA 98:5578–5583.

51. Nguyen AK, Molley TG, Kardia E, Ganda S, Chakraborty S, Wong SL, Ruan J, Yee BE, Mata J, Vijayan A, et al. (2023) Hierarchical assembly of tryptophan zipper peptides into stress-relaxing bioactive hydrogels. Nat. Commun. 14:6604.

52. Yamamura Y, Hirakawa H, Yamaguchi S, Nagamune T (2011) Enhancement of sortase A-mediated protein ligation by inducing a β-hairpin structure around the ligation site. Chem. Commun. 47:4742–4744.

53. Shao Q, Jiang Y, Yang ZJ (2022) EnzyHTP: A High-Throughput Computational Platform for Enzyme Modeling. J. Chem. Inf. Model. 62:647–655.

54. Jurich C, Shao Q, Ran X, Yang ZJ (2025) Physics-based modeling in the new era of enzyme engineering. Nat. Comput. Sci. 5:279–291.

55. Campbell BC, Paez-Segala MG, Looger LL, Petsko GA, Liu CF (2022) Chemically stable fluorescent proteins for advanced microscopy. Nat. Methods 19:1612–1621.

