## Supplemental Information for Wachsman et al. for "A FRET Ligation Assay using Fluorescent Proteins for Bacterial Sortase Enzymes"

**Table of Contents**

**Figure S1. Quantification of sortase-mediated ligation assay using SDS-PAGE and FIJI analyses.** 2

**Sequences used in study.** 3

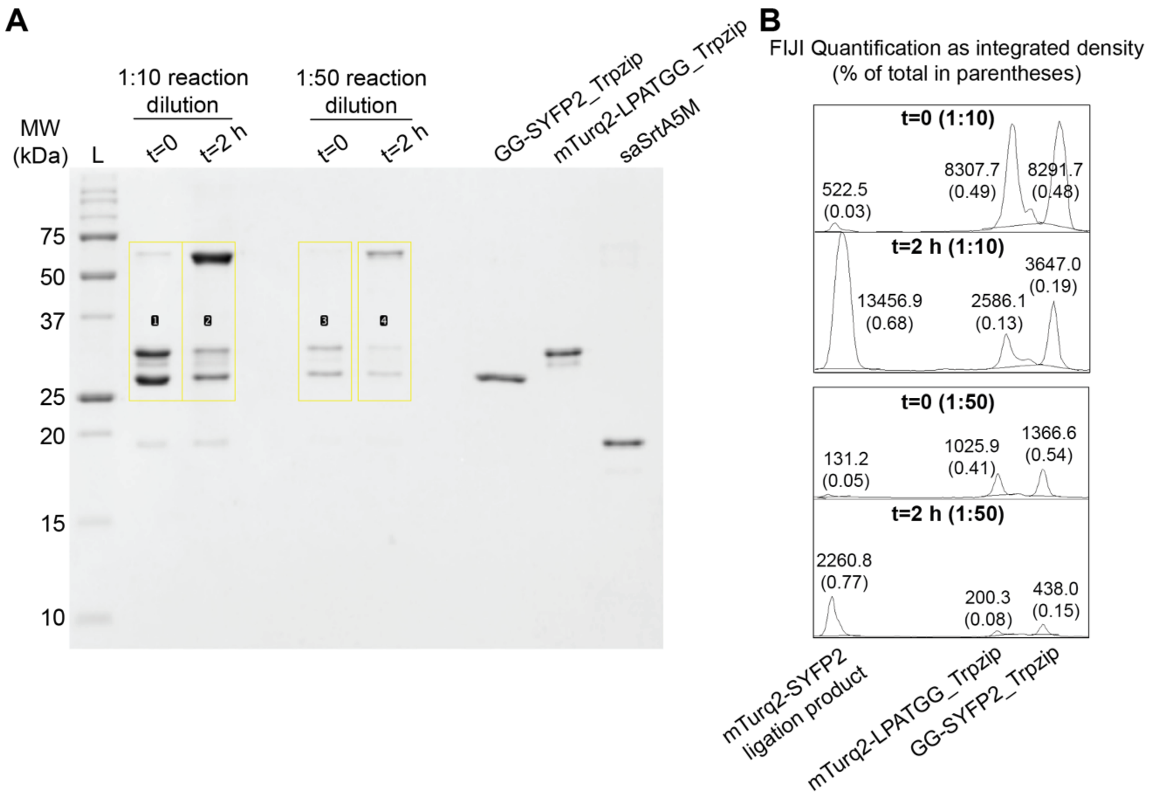


**Figure S1. Quantification of sortase-mediated ligation assay using SDS-PAGE and FIJI analyses.** (**A**) The full SDS-PAGE gel used for quantification. (**B**) The integrated density based on the peaks shown in (**A**) for both 1:10 dilutions (lanes 1-2) and 1:50 dilutions (lanes 3-4) of the quenched reaction samples. Total ligation efficiency was calculated as the density value of the ligation product (13456.9) / total density (13456.9 + 2586.1 + 3647.0) = 0.68 (or 68%) for the 1:10 reaction samples and 2260.8/(2260.8+200.3+438.0) = 0.77 (or 77%) for the 1:50 reaction samples.

**Sequences used in this study.** Underlined amino acids indicate the 6xHis tag used for protein purification and TEV protease cleavage site (ENLYFQ/S or ENLYFQ/G, where “/” is the cleavage site). **Bold and underlined** indicate linker sequences. **Bold** indicates the LPATG substrate recognition sequence. All sequences were inserted into the pET28a(+) expression plasmid. The wild-type saSrtA sequence matches UniProt ID SRTA_STAA8.

>mTurq2-LPATGG
MESSHHHHHHENLYFQSVSKGEELFTGVVPILVELDGDVNGHKFSVSGEGEGDATYGKLTLKFICTTGKLPVPWPTL
VTTLSWGVQCFARYPDHMKQHDFFKSAMPEGYVQERTIFFKDDGNYKTRAEVKFEGDTLVNRIELKGIDFKEDGNIL
GHKLEYNYFSDNVYITADKQKNGIKANFKIRHNIEDGGVQLADHYQQNTPIGDGPVLLPDNHYLSTQSKLSKDPNEK
RDHMVLLEFVTAAGITLGMDELYK**GGSLPATG**GGGG

> mTurq2-LPATGG_Trpzip
MESSHHHHHHENLYFQSVSKGEELFTGVVPILVELDGDVNGHKFSVSGEGEGDATYGKLTLKFICTTGKLPVPWPTL
VTTLSWGVQCFARYPDHMKQHDFFKSAMPEGYVQERTIFFKDDGNYKTRAEVKFEGDTLVNRIELKGIDFKEDGNIL
GHKLEYNYFSDNVYITADKQKNGIKANFKIRHNIEDGGVQLADHYQQNTPIGDGPVLLPDNHYLSTQSKLSKDPNEK
RDHMVLLEFVTAAGITLGMDELYK**GWTWTWGGSGGGGSLPATG**GGGG

>GG-SYFP2
MESSHHHHHHENLYFQG**GGGGSGGGGS**VSKGEELFTGVVPILVELDGDVNGHKFSVSGEGEGDATYGKLTLKLICTT
GKLPVPWPTLVTTLGYGVQCFARYPDHMKQHDFFKSAMPEGYVQERTIFFKDDGNYKTRAEVKFEGDTLVNRIELKG
IDFKEDGNILGHKLEYNYNSHNVYITADKQKNGIKANFKIRHNIEDGGVQLADHYQQNTPIGDGPVLLPDNHYLSYQ
SKLSKDPNEKRDHMVLLEFVTAAGITLGMDELYKGSGC

>GG-SYFP2_Trpzip
MESSHHHHHHENLYFQG**GGGWTWTWGS**VSKGEELFTGVVPILVELDGDVNGHKFSVSGEGEGDATYGKLTLKLICTT
GKLPVPWPTLVTTLGYGVQCFARYPDHMKQHDFFKSAMPEGYVQERTIFFKDDGNYKTRAEVKFEGDTLVNRIELKG
IDFKEDGNILGHKLEYNYNSHNVYITADKQKNGIKANFKIRHNIEDGGVQLADHYQQNTPIGDGPVLLPDNHYLSYQ
SKLSKDPNEKRDHMVLLEFVTAAGITLGMDELYKGSGC

>EGFP-LPETGG
MVSKGEELFTGVVPILVELDGDVNGHKFSVSGEGEGDATYGKLTLKFICTTGKLPVPWPTLVTTLTYGVQCFSRYPD
HMKQHDFFKSAMPEGYVQERTIFFKDDGNYKTRAEVKFEGDTLVNRIELKGIDFKEDGNILGHKLEYNYNSHNVYIM
ADKQKNGIKVNFKIRHNIEDGSVQLADHYQQNTPIGDGPVLLPDNHYLSTQSALSKDPNEKRDHMVLLEFVTAAGIT
LGMDELYK**LPETG**GLEHHHHHH

>GG-cpVenus
MESSHHHHHHENLYFQG**GG**MHEEEFQFLRCQQCQAEAKCPKLLPCLHTLCSGCLEASGMQCPICQAPWPLGADTPAL
ELMDGGVQLADHYQQNTPIGDGPVLLPDNHYLSYQSALSKDPNEKRDHMVLLEFVTAAGITLGMDELYKGGSGGMVS
KGEELFTGVVPILVELDGDVNGHKFSVSGEGEGDATYGKLTLKLICTTGKLPVPWPTLVTTLGYGLQCFARYPDHMK
QHDFFKSAMPEGYVQERTIFFKDDGNYKTRAEVKFEGDTLVNRIELKGIDFKEDGNILGHKLEYNYNSHNVYITADK
QKNGIKANFKIRHNIELE

>saSrtA5M
MESSHHHHHHENLYFQSQAKPQIPKDKSKVAGYIEIPDADIKEPVYPGPATREQLNRGVSFAEENESLDDQNISIAG
HTFIDRPNYQFTNLKAAKKGSMVYFKVGNETRKYKMTSIRNVKPTAVGVLDEQKGKDKQLTLITCDDYNEETGVWET
RKIFVATEVK
